# Parabrachial CGRP neurons modulates conditioned active defensive behavior under a naturalistic threat

**DOI:** 10.1101/2024.07.23.604768

**Authors:** Gyeong Hee Pyeon, Hyewon Cho, Byung Min Chung, June-Seek Choi, Yong Sang Jo

## Abstract

Recent studies suggest that calcitonin gene-related peptide (CGRP) neurons in the parabrachial nucleus (PBN) represent aversive information and signal a general alarm to the forebrain. If CGRP neurons serve as a true general alarm, activation of CGRP neurons can trigger either freezing or fleeing defensive behavior, depending on the circumstances. However, the majority of previous findings have reported that CGRP neurons modulate only freezing behavior. Thus, the present study examined the role of CGRP neurons in active defensive behavior, using a predator-like robot programmed to chase mice in fear conditioning. Our electrophysiological results showed that CGRP neurons encoded the intensity of various unconditioned stimuli (US) through different firing durations and amplitudes. Optogenetic and behavioral results revealed that activation of CGRP neurons in the presence of the chasing robot intensified fear memory and significantly elevated conditioned fleeing behavior during recall of an aversive memory. Animals with inactivated CGRP neurons exhibited significantly low levels of fleeing behavior even when the robot was set to be more threatening during conditioning. Our findings expand the known role of CGRP neurons in the PBN as a crucial part of the brain’s alarm system, showing they can regulate not only passive but also active defensive behaviors.

## Introduction

Effective survival necessitates a repertoire of dynamic defensive behaviors, encompassing both passive and active responses. Passive defensive strategies, such as freezing, help avoid detection from predators by reducing motion (Blanchard & Blanchard, 1969a; Fanselow, 1980, 1982). In contrast, active defensive behaviors, including fleeing or fighting, enable animals to swiftly escape or confront imminent threats (Blanchard & Blanchard, 1969b; Bolles, 1970). The ability to adaptively switch between passive and active defenses in response to varying threat contexts is essential for optimizing survival outcomes, as demonstrated by studies utilizing naturalistic threat stimuli like predator-like robots or looming disks, which allowed the observation of various critical defensive behaviors (Choi & Kim, 2010; Kang et al., 2022; Pyeon et al., 2023; Telensky et al., 2011). A critical component of this adaptive response is the general alarm signal, which triggers appropriate defensive behaviors in the face of danger. These signals help organisms quickly recognize and respond to potential threats. The mechanisms underlying these alarm signals can be studied through Pavlovian fear conditioning (Bolles & Collier, 1976; Fanselow & Poulos, 2005; LeDoux, 2000; Maren, 2001). In this process, a neutral sensory stimulus (conditioned stimulus or CS) is paired with an aversive unconditioned stimulus (US), leading to a conditioned response (CR) that can be expressed as either freezing or fleeing, depending on the specific features of the CS (Borkar & Fadok, 2021; Fadok et al., 2017) and US (Lee et al., 2018; Pyeon et al., 2023).

Neurons within the parabrachial nucleus (PBN) that express calcitonin gene-related peptide (CGRP) have been suggested to function as general alarm signals in the brain (Palmiter, 2018). These neurons respond to noxious stimuli of diverse sensory modalities (Campos et al., 2018; Carter et al., 2013; Chen et al., 2018; Kang et al., 2022) and transmit interoceptive and exteroceptive information to the forebrain (Bernard & Besson, 1988; Chiang et al., 2019).

Additionally, these CGRP neurons relay US information to the central amygdala during conventional fear conditioning with electric footshock (Han et al., 2015). Prior studies have primarily focused on the role of CGRP neurons in mediating passive freezing behavior, demonstrating that activation of these neurons exclusively elicits immediate freezing behavior and contributes to the formation of fear memories (Bowen et al., 2020; Han et al., 2015). However, for CGRP neurons to function as true general alarm system, they must capable of trigerring both passive and active defensive behaviors, depending on the severity, immediacy, or type of threat. While the role of CGRP neurons in passive responses is well-established, their potential involvement in active defensive behaviors remains unexplored.

To address this, we employed a more dynamic and ecologically relevant US by using a predator-like robot to chase the animals, thereby incorporating an imminent threat. We hypothesized that CGRP neurons contribute to the selection of adaptive defensive behaviors by signaling the severity of the threat. We first recorded CGRP neuron activity in response to various aversive stimuli including the robot chasing to determine whether they encode noxious stimuli differentially. We then manipulated CGRP activity—both activating and inactivating—during fear conditioning with robot chasing and footshock. Our results demonstrate that manipulation of CGRP neurons bidirectionally modulates conditioned fleeing behaviors through altering the perception of the threat. These results support the role of CGRP neurons as a general alarm system, capable of inducing both passive and active defensive behaviors.

## Results

### Differential responses of CGRP neurons to aversive stimuli of varying intensities

The response profiles of CGRP neurons in conventional fear conditioning with footshock have been well reported (Han et al., 2015). However, how CGRP neurons respond to chasing threat has not been established. To investigate the activity of CGRP neurons in response to robot chasing, *in vivo* recordings using the optical-tagging strategy were performed (Jo et al., 2018; Juarez et al., 2023). Heterozygous mice expressing Cre-recombinase at the *Calca* locus (*Calca^Cre^*^/+^) were injected with a Cre-dependent adeno-associated virus (AAV) carrying an excitatory channelrhodopsin (ChR2) with red fluorescent protein (AAV-DIO-ChR2-mScarlet). Then, a movable optrode array containing one optic fiber with four tetrodes was implanted over the PBN (Figure 1A).

**Figure 1.**
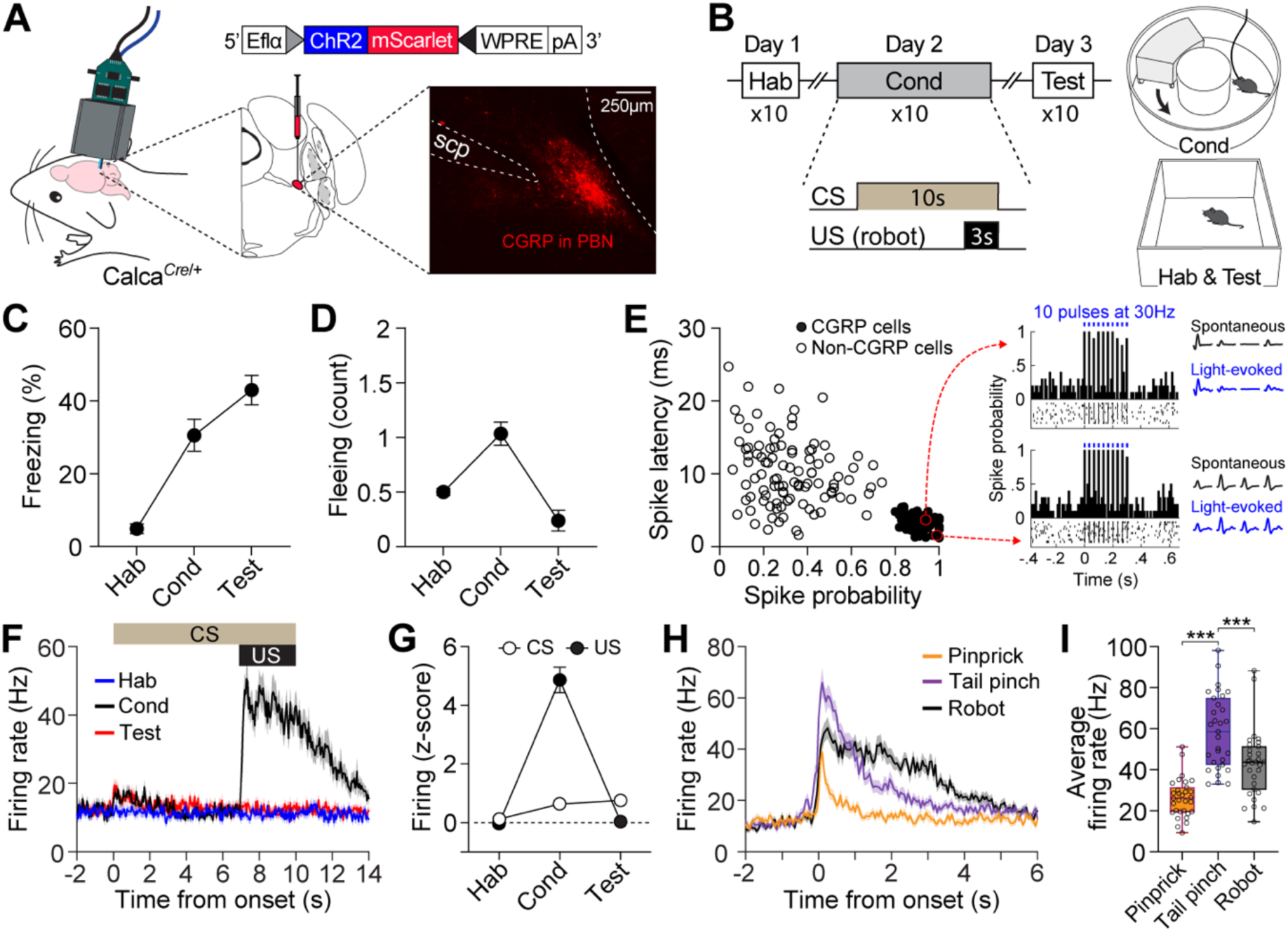
Distinct firing response patterns of CGRP neurons to different aversive stimuli. **(A)** Schematic of AAV-DIO-ChR2-mScarlet injection and optrode implantation into the PBN of *Calca^Cre^*^/+^ mice (n = 6) and the corresponding representative histology image. **(B)** Procedures for fear conditioning experiments with the chasing robot and a schematic diagram of the context used. **(C-D)** Freezing (C) and fleeing behaviors (D) in response to the CS during habituation, conditioning, and retention test. **(E)** Characteristics of light-evoked responses. Neurons with a short spike latency and a high spike probability response to light stimulation (filled circles) were classified as CGRP neurons. Inset: histograms showing firing patterns of two representative opto-tagged CGRP neurons response to 10 blue light pulses at 30 Hz. **(F)** Population firing rates of all recorded CGRP neurons (hab: n = 28; cond: n = 29, test: n = 27) during fear conditioning with the robot. (**G**) Normalized firing in response to CS and US. (**H**) Population responses of all recorded CGRP neurons (n = 31) in response to three aversive stimuli. (**I**) Average firing rates of CGRP neurons to pinprick, tail pinch, and robot chasing. Among these aversive stimuli, tail pinch induced a significantly greater increase in firing rates compared to both pinprick and robot chasing (one-way ANOVA, F(2, 90) = 35.87, *p* < 0.001; post-hoc tests, *p* values < 0.001). \*\*\**p* < 0.001.

After 2 weeks of recovery, neuronal activity was recorded during fear conditioning with a robot over 3 consecutive days (Figure 1B). Animals were first habituated to a tone (4 kHz; 70 dB; 10 s) as a CS in a rectangular box. The following day, the animals were placed in a donut-shaped maze and presented with the CS 10 times, each paired with an US of being chased by the robot at a speed of 70 cm/s for 3 s. On day 3, fear memory was assessed by presenting the CS alone 10 times in the same context as the habituation session. In this behavioral paradigm, animals exhibited both freezing and fleeing responses to the CS during conditioning (Figure 1C and D). However, on the test day, when the robot was no longer presented, fleeing responses were no longer observed, and the animals showed increased freezing.

To identify CGRP neurons, 10 pulses of blue light (5-ms duration) at 30 Hz were delivered 10 times at the end of each behavioral recording session. Out of 183 PBN neurons, 84 cells with a high probability of light-evoked spikes (> 0.8) and a short spike latency (< 5.5 ms) after light onset were classified as CGRP neurons (Figure 1E). Compared to habituation, CGRP neurons showed significantly increased excitation to the CS during conditioning and retention, but only within the first 1 s after CS onset (1.5-fold increase); this difference became non-significant starting at 2 s (Figure 1G). However, these neurons exhibited significant excitation to the US with a 4-fold increase (Figure 1F and G). Our findings using the robot as the US revealed that CGRP neurons primarily represent US information, albeit to a lesser extent, the onset of US-predictive information.

Given that CGRP neurons preferentially respond to aversive US, we next asked how CGRP neurons encode different types of aversive stimuli. To address this, we monitored the activity of CGRP neurons while the animals received three types of stimuli, each varying in perceived threat intensity: 1) a pinprick to the hind paw using a needle (approximately 0.5 s); 2) a tail pinch 2 cm from the tail base using forceps (1 s); and 3) being chased by a robot (3 s). These aversive stimuli elicited different defensive behaviors. The pinprick caused hind paw withdrawal, the tail pinch triggered vocalizations (audible squeaks) and immediate escaping behavior, indicating the highest threat intensity, while the robot chasing prompted only escaping behavior without any vocalization. CGRP neurons showed significantly excited firing that was time-locked to the onset of all three aversive stimuli and maintained this activity throughout the duration of each stimulus. After the offset of the stimuli, neuronal activity gradually restored back to the baseline with a slight delay (Figure 1H). These aversive stimuli elicited significantly different amplitudes of firing rates (Figure 1I). During the tail pinch, which generated the strongest defensive behavior, CGRP neurons exhibited the highest excitation amplitude, ranging from 33 to 98 Hz, with an average of 58 Hz. Taken together, these results demonstrate that CGRP neurons represent the temporal characteristics and intensity of different aversive stimuli through variations in firing duration and amplitude.

### CGRP activation promotes conditioned fleeing in robot chasing and conditioned freezing in footshock

To determine whether increasing or decreasing CGRP neuronal activity would induce defensive behaviors other than freezing, we first observed which defensive behaviors were elicited by either stimulating or inhibiting CGRP neurons in the absence of any external stimuli. *Calca^Cre^*^/+^ mice were randomly assigned to groups and bilaterally injected with either AAV-DIO-ChR2-mScarlet for activation, AAV-DIO-Jaws-GFP for inactivation, or AAV-DIO-eYFP for control, followed by the implantation of optic fibers over the PBN (Figure 2A). To activate CGRP neurons, mice received 30 s of 40 Hz photostimulation, delivered 4 times, based on the observed spontaneous firing rate of approximately 43 Hz in response to robot chasing (Figure 1I). For inactivation, CGRP neurons were inhibited for 2 s, followed by a 1-s ramp down, repeated in cycles until a total duration of 30 s. Jaws and control groups showed no difference in movement during light on and off phases, indicating that light delivery did not alter their defensive behavior. However, activation of CGRP neurons immediately induced robust freezing behavior, consistent with previous studies (Bowen et al., 2020; Han et al., 2015). These results confirmed that stimulating CGRP neurons without external aversive stimuli generates rapid, unconditioned freezing behavior in mice.

**Figure 2.**
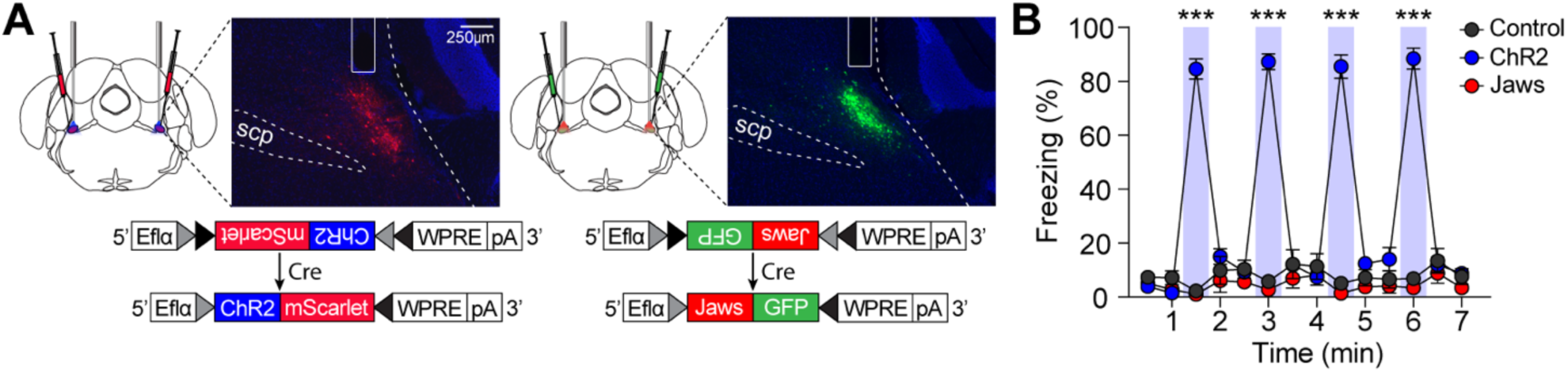
Stimulation of CGRP neurons in the absence of any external stimuli induces robust freezing behavior. **(A)** Schematic of bilateral AAV-DIO-ChR2-mScarlet or AAV-DIO-Jaws-GFP injections and optic fiber implantation into the PBN, with representative histological images of viral expression. **(B)** 30 s stimulation of CGRP neurons at 40 Hz resulted in significantly higher time-locked freezing behaviors in the ChR2 group compared to both the Jaws and control groups (n = 10 per group; significant group × time interaction in a mixed-design ANOVA, F(26, 351) = 61.32, *p* < 0.001; post-hoc tests at each time, *p* values < 0.001). \*\*\**p* < 0.001.

We then tested whether manipulating activity of CGRP neurons during fear conditioning with robot chasing promotes fleeing behavior or amplifies freezing behavior. To adequately increase the activation of CGRP neurons within a physiologically relevant range while animals were being chased by the robot, we optogenetically stimulated CGRP neurons at 30 Hz. Combined with the spontaneous firing rate during robot chasing (about 40 Hz), this totaled approximately 70 Hz, matching the peak firing rate observed in the upper quartile during tail pinch (Figure 1I). *Calca^Cre^*^/+^ mice underwent the fear conditioning paradigm in which the CS was paired with the robot chasing (3 s, 70 cm/s), with CGRP neurons selectively activated (30 Hz) or inhibited (3 s on and 1-s ramp down) throughout the presentation of the chasing (Figure 3A). Physical bumping occurred during robot chasing, influencing perceived threat, with more bumps leading to increased fear. We found no significant differences in bumping incidents across the three groups, suggesting that differences observed later were due to CGRP neuronal activity regulation (Figure 3−figure supplement 1C).

**Figure 3.**
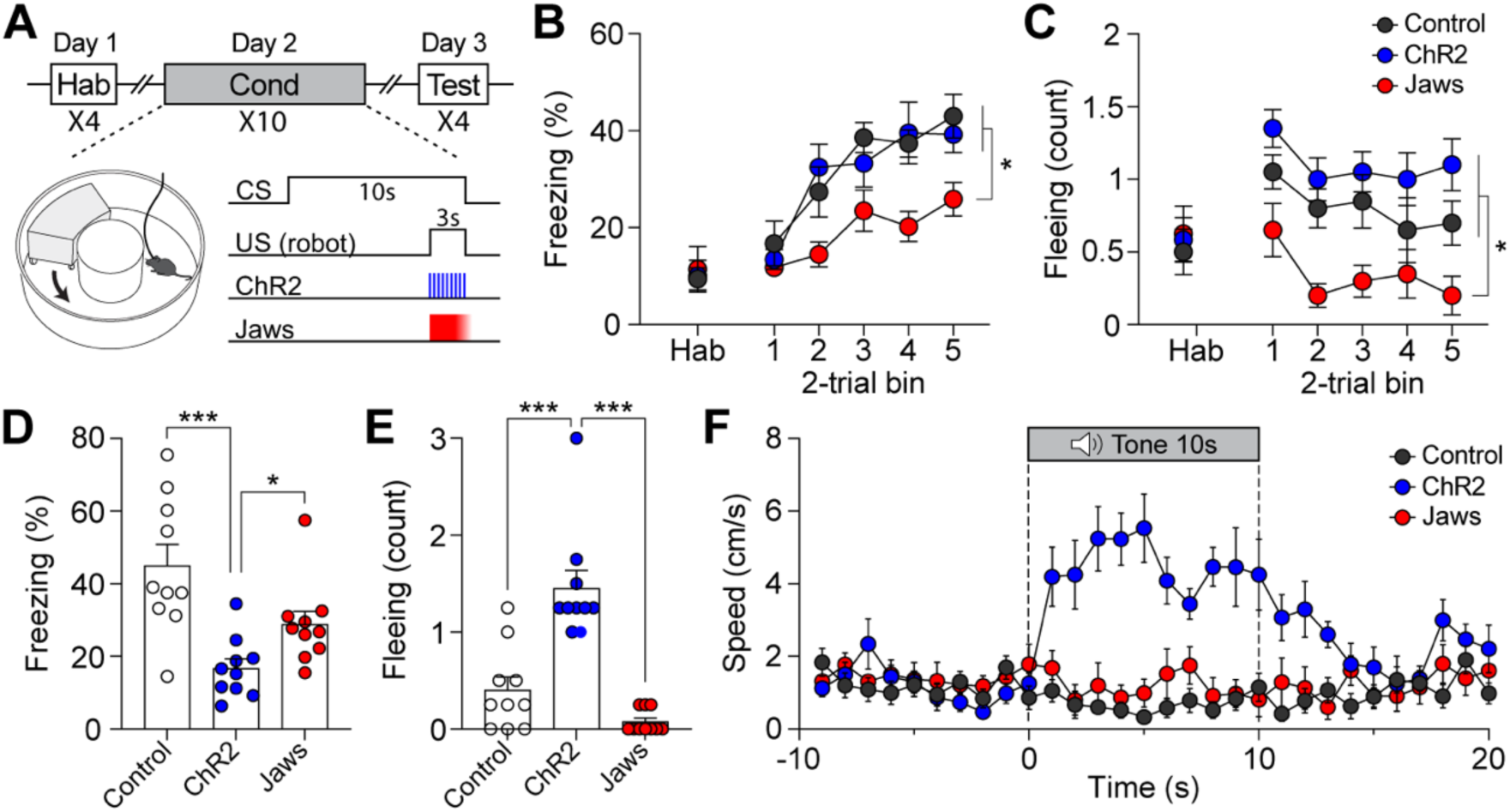
Activation of CGRP neurons enhances active defensive behavior. **(A)** A schematic diagram of fear conditioning protocol with the robot. CGRP neuronal activity was bidirectionally manipulated during the presentation of the robot chasing. **(B)** Freezing to the CS during habituation and conditioning sessions. A progressive increase in freezing was observed in all three groups (n = 10 per group), but the Jaws group showed significantly lower freezing levels compared to the other two groups (significant group effect in a mixed-design ANOVA, F(2, 27) = 6.99, *p* < 0.01; subsequent post-hoc tests, *p* values < 0.01). **(C)** Fleeing in response to the CS during habituation and conditioning sessions. Both the ChR2 and control groups displayed equivalent levels of fleeing response, while the Jaws group showed lower levels of fleeing compared to the other two groups (significant group effect in a mixed-design ANOVA, F(2, 27) = 12.60, *p* < 0.001; post-hoc tests, *p* values < 0.05). **(D)** Average freezing in response to the CS during the retention test. Both the ChR2 and Jaws groups froze significantly less than the control group (one-way ANOVA, F(2, 27) = 11.01, *p* < 0.001; post-hoc test, *p* values < 0.05). **(E)** Average fleeing to the CS during the retention test. The ChR2 group showed significantly more fleeing behaviors compared to both the Jaws and control groups (one-way ANOVA, F(2, 27) = 28.57, *p* < 0.001; subsequent post-hoc test, *p* values < 0.001). **(F)** Average velocities in response to the CS during the retention test. The average velocity of the ChR2 group during the CS was significantly higher compared to that observed in the Jaws and control groups (significant group effect in a mixed-design ANOVA, F(2, 27) = 58.60, *p* < 0.001; post-hoc test, *p* values < 0.01). \**p* < 0.05, \*\*\**p* < 0.001.

During conditioning, both ChR2 and control groups showed a progressive increase in freezing, reaching comparable levels (Figure 3B). In contrast, the Jaws group exhibited significantly lower freezing levels than the other two groups. In terms of fleeing responses on the conditioning day, both ChR2 and control mice displayed significantly higher levels of fleeing to the CS, whereas the Jaws group had a consistently low fleeing response (Figure 3C). Fear memory was assessed 24 h after conditioning by presenting four CSs alone. During the retention test, the ChR2 group exhibited fleeing as their primary defensive behavior instead of freezing, whereas the control group showed dominant freezing behavior (Figure 3D and E; Movie 1). Additionally, analysis of movement velocity revealed that ChR2 mice had a higher fleeing speed compared to the other two groups (Figure 3F). We expected the Jaws group to show lower freezing and fleeing responses due to inhibited US signaling during conditioning. However, they exhibited similar freezing levels to the control group (Figure 3D). Considering that CGRP neurons continued to fire even after the aversive stimuli ceased (Figure 1F), the Jaws group was able to form fear memory due to CGRP activation during the prolonged period following the US. Alternatively, the Jaws group acquired fear memory through US information processed via other pathway distinct from CGRP neurons (Johansen et al., 2010; Kang et al., 2022). Taken together, these findings demonstrate that enhanced CGRP activity during imminent threat stimuli strengthens fear memory, causing fleeing responses observed during conditioning to persist through the retention test.

We next examined whether the same modulation of CGRP neuron activity paired with electric footshock (1s; 0.3 mA) would also engage in active defensive behavior. After a one-week of resting period following the conditioning paradigm with the robot, the same group of mice underwent conventional fear conditioning (Figure 4A). Although the CS used in the conventional fear conditioning (12 kHz; 70 dB; 10 s) differed from the one used with the robot (4 kHz; 70 dB; 10 s), residual effects were observed in the ChR2 group during the habituation session. The ChR2 group exhibited similarly minimal levels of freezing compared to the control group but demonstrated significantly higher fleeing behavior (Figure 4C).

**Figure 4.**
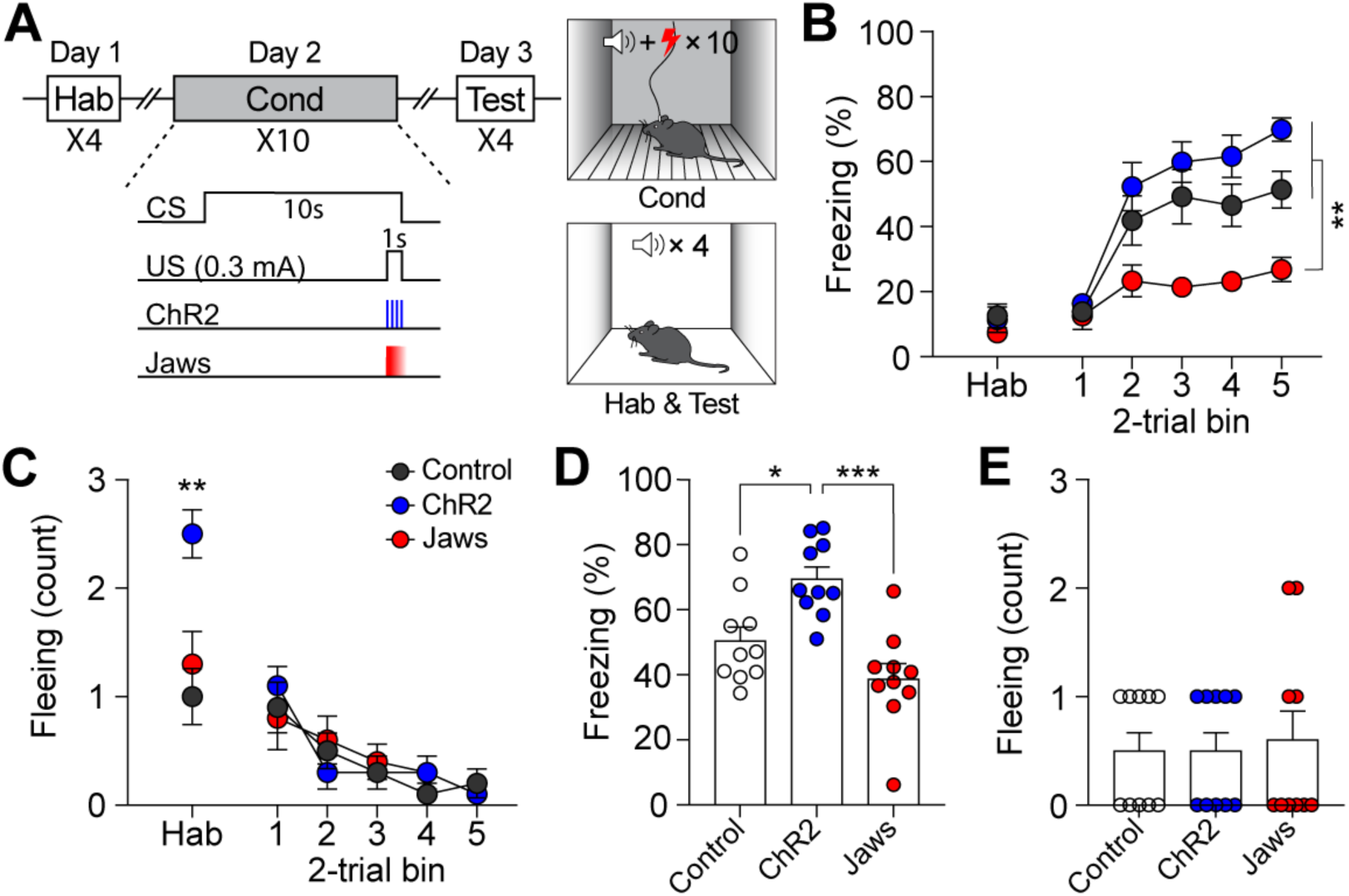
CGRP activation during conventional fear conditioning promotes passive, but not active, defensive behavior. **(A)** A schematic diagram of fear conditioning protocol with the footshock. **(B)** Freezing to the CS during habituation and conditioning sessions. All three groups showed a progressive increase in freezing as trials progressed, but the Jaws group froze significantly less compared to the other two groups (significant group effect in a mixed-design ANOVA, F(2, 27) = 19.74, *p* < 0.001; subsequent post-hoc tests, *p* values < 0.01). **(C)** Fleeing in response to the CS during habituation and conditioning sessions. The ChR2 group showed significantly higher fleeing responses during habituation, suggesting some residual effect from fear conditioning with the robot (one-way ANOVA, F(2, 27) = 9.8, *p* < 0.01; post-hoc test, *p* values < 0.01). All three groups displayed a significant decrease in fleeing behavior during conditioning, with no group differences observed (no group effect in a mixed-design ANOVA, F(2, 27) = 0.07, *p* = .93). **(D)** Average freezing in response to the CS during the retention test. The ChR2 group froze significantly more than the Jaws and control group (one-way ANOVA, F(2, 27) = 13.31, *p* < 0.001; post-hoc tests, *p* values < 0.05). **(E)** Average fleeing to the CS during the retention test. Fleeing responses were minimal across all three groups, and no significant differences were observed (one-way ANOVA, F(2, 27) = 0.08, *p* = 0.92). \**p* < 0.05, ** *p* < 0.01, \*\*\**p* < 0.001.

During conditioning, both ChR2 and control groups exhibited a gradual increase in freezing as the trials progressed (Figure 4B). However, consistent with the previous experiment with the robot, the Jaws group showed significantly lower levels of freezing compared to the other two groups. Moreover, all three groups showed equivalently low levels of fleeing responses (Figure 4C). When fear memory was tested 24 h later, ChR2-expressing mice displayed significantly more freezing compared to both control and Jaws-expressing mice (Figure 4E). The Jaws and control groups exhibited similar levels of freezing, with no significant difference between the two groups. In terms of fleeing response, since all three groups demonstrated minimal fleeing responses, there was no significant difference observed (Figure 4F). These data show that the same CGRP stimulation did not promote fleeing responses; however, with footshock as the US, the freezing response observed during conditioning was intensified in the retention test. Overall, additional activation of CGRP neurons enhances fear memory, resulting in conditioned fleeing responses following robot chasing and conditioned freezing responses after footshock.

### CGRP neurons intensify threat perceptions and regulate defensive behaviors

To further investigate whether the previously observed fleeing responses with additional CGRP activation in the presence of the robot (Figure 3E) were due to the intensified perception of the US threat, we systematically escalated the threat level of the US by increasing the robot’s speed without manipulating CGRP activity. In the previous experiment, the robot moved at a speed of 70 cm/s, making one and a half turns in the donut maze within 3 seconds. By increasing the speed to 80 cm/s, the robot made two full turns, and at 90 cm/s, it made two and a half turns within the same time frame. Additionally, we analyzed the correlation between robot speed and the number of physical bumps, revealing a significant positive relationship in which higher robot speeds led to more bumps (Figure 5−figure supplement 1A and B). These findings suggest that the increased robot speed resulted in the animals perceiving a greater threat due to more physical contact.

On the conditioning day, all three groups showed equivalent levels of freezing and fleeing behavior (Figure 5A and B). However, on the retention test day, mice exposed to speeds of 70 cm/s and 80 cm/s showed no group differences in freezing, while those subjected to 90 cm/s exhibited significantly lower levels of freezing compared to the other two groups (Figure 5C). In contrast, animals in the 90 cm/s speed displayed a significantly higher number of fleeing responses compared to the other two groups (Figure 5D and E). There was a positive correlation between robot speed and fleeing responses, and a negative correlation between robot speed and freezing responses (Figure 5C and D). Furthermore, animals exposed to the 90 cm/s speed exhibited a fleeing response similar to that of those subjected to 70 cm/s robot chasing with CGRP stimulation (Figure 3E). These findings confirm that the increased activity of CGRP neurons enhances fleeing behavior by intensifying the perceived threat of the US.

**Figure 5.**
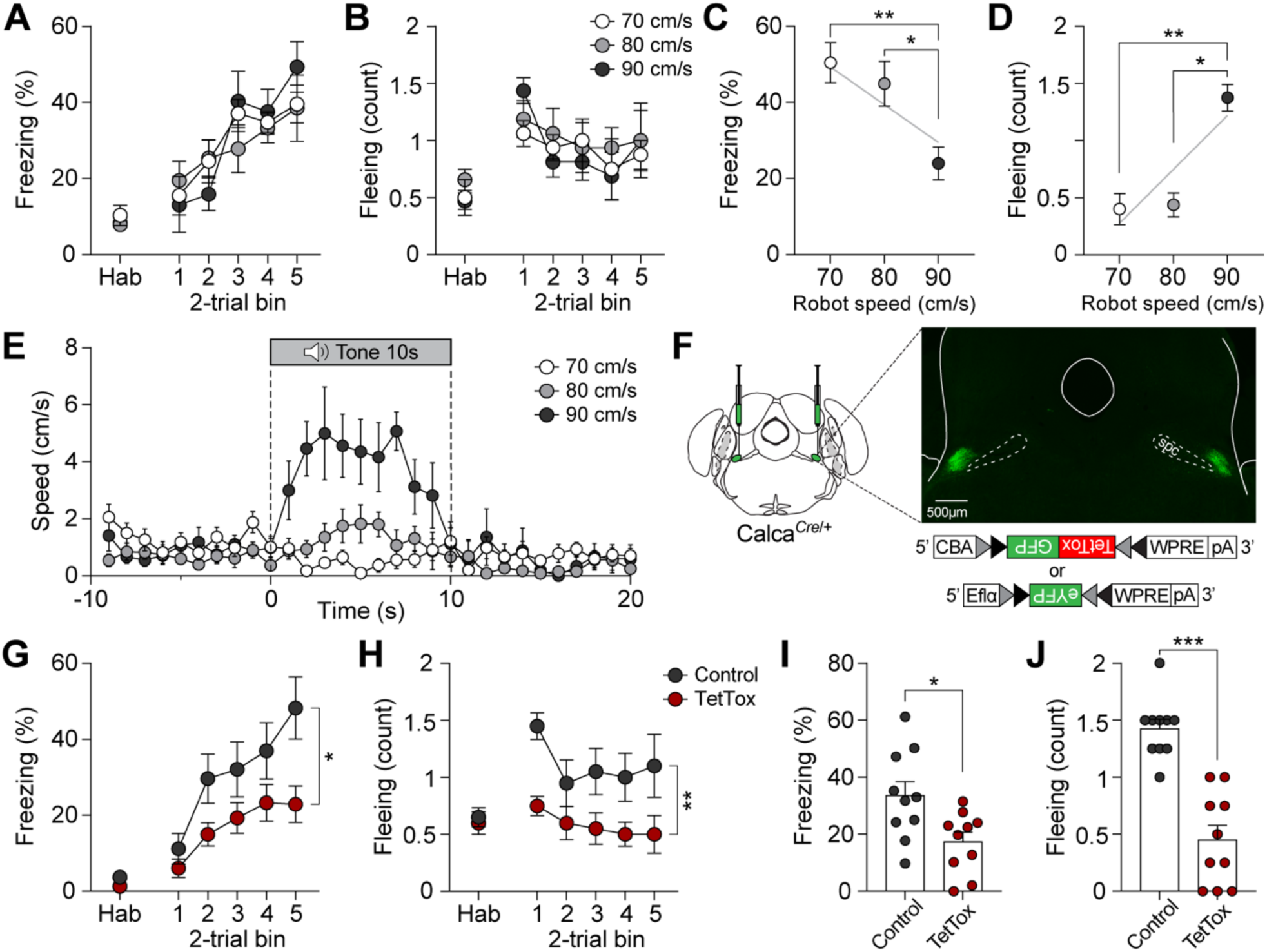
CGRP neurons modulate defensive behavior by enhancing the intensity of threats. **(A)** Freezing to the CS during habituation and conditioning sessions for groups subjected to three different robot speeds (n = 8 per group). All three groups showed an equivalent progressive increase in freezing as trials progressed, with no significant differences between the groups (mixed-design ANOVA, F(2, 21) = 0.11, *p* = 0.89). **(B)** Fleeing in response to the CS during habituation and conditioning sessions. Fleeing responses were similar across all three groups (no group effect in a mixed-design ANOVA, F(2, 21) = 0.22, *p* = 0.81). **(C)** Average freezing in response to the CS during the retention test. Animals exposed to 70 or 80 cm/s robot speed froze significantly more compared to those subjected to 90 cm/s (one-way ANOVA, F(2, 21) = 6.60, *p* < 0.01; post-hoc tests, *p* values < 0.05). There was a negative correlation between freezing responses and robot speed (gray line; *r* = -0.61, *p* < 0.01). **(D)** Average fleeing to the CS during the retention test. Animals subjected to 90 cm/s robot speed displayed significantly more fleeing responses compared to both 70 and 80 cm/s (one-way ANOVA, F(2, 21) = 17.18, *p* < 0.001; subsequent post-hoc tests, *p* values < 0.001). Fleeing responses were positively correlated with robot speed (gray line; *r* = 0.67, *p* < 0.001). **(E)** Average velocities in response to the CS during the retention test. The average velocity of the 90 cm/s group during the CS was significantly higher than that observed in the 70 and 80 cm/s groups (significant group effect in a mixed-design ANOVA, F(2, 27) = 58.60, *p* < 0.001; post-hoc test, *p* values < 0.01). **(F)** Schematic of bilateral AAV-DIO-TetTox-GFP or AAV-DIO-eYFP (control) injections into the PBN and representative histological images of TetTox expression. **(G)** Freezing to the CS during habituation and conditioning sessions (n = 10 per group). Both groups showed a progressive increase in freezing, but the TetTox group, with inactivated CGRP neurons, exhibited significantly lower levels of freezing compared to the control group (significant group effect in a mixed-design ANOVA, F(1, 18) = 6.42, *p* < 0.05). **(H)** Fleeing in response to the CS during habituation and conditioning sessions. The control group showed significantly greater levels of fleeing responses compared to the TetTox group (significant group effect in a mixed-design ANOVA, F(1, 18) = 12.74, *p* < 0.01). **(I)** Average freezing to the CS during the retention test. The TetTox group displayed significantly lower levels of freezing compared to the control group (independent t-test, t(18) = 2.7, *p* < 0.05). **(J)** Average fleeing in response to the CS during the retention test. The TetTox group showed significantly lower levels of fleeing compared to the control group (independent t-test, t(18) = 6.37, *p* < 0.001). \**p* < 0.05, ** *p* < 0.01, \*\*\**p* < 0.001.

We next sought to confirm whether CGRP neurons are necessary for inducing active defensive behavior under high-speed conditions. Since Jaws inhibition was insufficient to block fear memory formation (Figure 3D), we bilaterally injected either Cre-dependent tetanus toxin light chain (TetTox; AAV-DIO-GFP) for effective silencing by selectively blocking neurotransmitter release, or AAV-DIO-eYFP (control) into the PBN. Mice then underwent fear conditioning with the robot at a speed of 90 cm/s (Figure 5F). On the conditioning day, the TetTox group exhibited significantly lower levels of both freezing and fleeing compared to the control group (Figure 5G and H). This persisted on the retention day, with the TetTox group consistently showing reduced levels of freezing and fleeing compared to controls (Figure 5I and J). These results suggest that CGRP neurons are necessary for perceiving threats and promoting fleeing. Enhancing CGRP neuronal activity, either optogenetically or by increasing threat levels, strengthens fear memory, leading to intensified active defensive behaviors during the retention test.

## Discussion

CGRP neurons, known for relaying US information to the forebrain and inducing passive freezing behavior (Han et al., 2015), were examined to explore their role in active defensive responses. Using a naturalistic paradigm with a robot and other aversive stimuli of varying threat levels, we recorded neuronal activity and found that CGRP neurons encode different threat intensities through variations in firing duration and amplitude. Enhancing these neurons during fear conditioning elevated percieved threat and reinforced fear memory, while inhibiting them weakened it. Heightened fear memory was expressed differently depending on the type of US: during robot chasing, enhanced CGRP activity promoted persistent fleeing in the retention test, while during footshock, it intensified freezing responses. Systematically escalating robot speed to increase threat intensity confirmed that higher threat levels strengthened conditioned fleeing responses, while silencing these neurons prevented the formation of conditioned active defensive behaviors. Overall, our results reveal that CGRP neurons not only elicit passive freezing but also engage in active defensive behaviors, establishing them as a comprehensive alarm system.

The choice of defensive strategies depends on the percieved severity of the threat (Fanselow & Lester, 2013). For instance, when an animal detects a predator at a relatively safe distance, freezing is the most likely defensive behavior, as it helps avoid detection. However, as the threat becomes more imminent and threat levels increase, freezing is no longer the optimal choice. At this point, the animal shifts from passive freezing to more active defenese, adopting behaviors such as fleeing or, if necessary, fighting. However, most research on CGRP has utilized footshock (Bowen et al., 2020; Han et al., 2015) or other various stimuli in small arenas (Kang et al., 2022), potentially limiting the observation of fleeing responses and leading to a focus on passive freezing as the predominant behavior studied. Thus, in our study, we introduced different types of US presenting an imminent threat. The predator-like robot allowed the animals to perceive the impending threat through dynamic sensory inputs, with the distance to the threat being determinable. Using this paradigm, we observd fleeing behavior in the animals, demonstrating that CGRP neurons are not only involved in passive freezing but also play a crucial role in acitive defensive behaviors.

Recent studies have modified conventional fear conditioning protocols to investigate active defensive behaviors in animals (Fadok et al., 2017). One such example is changing the CS to a serial-compound stimulus, where a pure tone is immediately followed by a white noise, inducing freezing and flight responses, respectively. While effective for observing transitions between freezing and fleeing, our electrophysiological data show that CGRP neurons are more excited in response to the US compared to the CS (Figure 1F and G). Thus, altering the type of US is more appropriate for stuyding CGRP neurons. In addition, the robot allowed us to systematically increase threat levels by adjusting its speed, providing a more controlled approach to studying defensive behaviors. Our results showed that a robot speed of 70 cm/s did not induce fleeing response during the retention test (Figure 5D); however, increasing the robot’s speed to 90 cm/s elicited conditioned flight responses in control mice. Moreover, CGRP activation combined with a 70 cm/s robot speed induced flight responses similar to those observed with a 90 cm/s robot speed in control mice. This demonstrates that CGRP activation amplifies the perceived threat, thereby promoting active defensive behaviors.

We inhibited CGRP neurons optogenetically in animals while being chased by the robot during conditioning. During this session, the Jaws group showed less freezing and fleeing compared to the control group. However, the reduced fear responses were not sustained on the retention test day, suggesting that transient inhibition during the chasing was insufficient to suppress fear memory formation (Figure 3D). Our electrophysiological results showed that after the 3-s chasing, CGRP neurons took about 2 to 3 s to return to baseline (Figure 1F). While CGRP neurons were sufficiently inhibited during the first 3 s, they may have become excited during the subsequent 2 to 3 s, presumably due to the pain experienced after bumping into the robot. This delayed excitation of CGRP neurons could have contributed to the formation of fear memory. In a subsequent experiment with increased robot speed, we used TetTox to silence CGRP neurons more effectively compared to temporary inhibition. This group consistently showed lower fear responses compared to the control group even on retention day, indicating a more significant impact on fear memory formation compared to transient inhibition. However, fear memory formation was still not completely blocked, suggesting the involvement of other pathways in processing the aversive stimuli. For instance, different populations of CGRP neurons in the parvocellular subparafascicular nucleus of the thalamus also respond to threats and relay negative emotional signals to the amygdala thereby contributing to aversive memory formation (Kang et al., 2022). Additionally, the midbrain periaqueductal gray (PAG) transmits aversive signals to the amygdala (Johansen et al., 2010; Johansen et al., 2012; Ozawa et al., 2017) and other forebrain structures (Lefler et al., 2020; Masferrer et al., 2020), ensuring effective expression of defensive responses upon the detection of a threat. Although memory formation can occur through various pathways, our use of optogenetics and TetTox demonstrated that CGRP neurons are critically involved in active defensive behavior.

In conclusion, by using both conventional footshock and a nauturalstic paradigm, the present study emphasizes the role of CGRP neurons in actively promoting both passive and active defensive behaviors. Futhermore, by observing the flight response during the recall of an aversive memory, we reveal that enhanced CGRP neuron activity alters threat perception, making animals respond as if a predator were chasing them at a faster speed. Together, this highlights their function as a comprehensive general alarm system, modulating appropriate defensive behaviors and ensuring animals effectively execute strategies to counter diverse and escalating dangers.

## Materials and methods

### Key resources table

| Reagent type (species) or resource | Designation | Source or reference | Identifiers | Additional information |
| --- | --- | --- | --- | --- |
| Strain, strain background (*Mus musculus*) | B6.Cg-Calca<sup>tm1.1(cre/EGFP)Rpa/J</sup> | The Jackson Laboratory | RRID:IMSR_JAX:033168 |  |
| Strain, strain background (C57BL/6J, C57BL/6J) | C57BL/6J | The Jackson Laboratory | RRID:IMSR_JAX:000664 |  |
| Recombinant DNA reagent | AAV-DIO-ChR2-mScarlet | IBS Virus Facility | N/A |  |
| Recombinant DNA reagent | AAV-DIO-Jaws-GFP | (Jo et al., 2018) | N/A |  |
| Recombinant DNA reagent | AAV-DIO-eYFP | Addgene | RRID:Addgene_27056 |  |
| Recombinant DNA reagent | AAV-DIO-TetTox-GFP | (Han et al., 2015) | N/A |  |
| Software, algorithm | ANY-maze 5.3 | Stoelting Co | N/A |  |
| Software, algorithm | GraphPad Prism | GraphPad software | N/A |  |
| Software, algorithm | SPSS | IBM | N/A |  |
| Software, algorithm | Offline Sorter | Plexon Inc | N/A |  |

### Animals

We used heterozygous *Calca^Cre^*^/+^ mice, generated by breeding Calca*^Cre/Cre^* (Cat. 033168) with C57BL/6J (Cat. 000664) from Jackson Laboratory. Both male and female mice, aged 3 to 6 months, were used in all studies, and no sex differences were observed. Mice were housed in a temperature- and humidity-controlled facility on a 12 h light/dark cycle (lights off at 7 AM) with ad libitum access to food and water. All experiments were performed during the dark phase of the cycle under the guidelines of the Institutional Animal Care and Use Committee at the Korea University (KUIACUC-2022-0057).

### Virus production

All AAV vectors were prepared as described previously (Pyeon et al., 2024). Cre-dependent optogenetic viruses included AAV-DIO-ChR2-mScarlet, AAV-DIO-Jaws-GFP, and AAV-DIO-eYFP (control). For selective inactivation of CGRP neurons, AAV-DIO-TetTox-GFP was used. Viral aliquots were stored at -80℃ before stereotaxic injection.

### Stereotaxic surgery

Mice were anesthetized with isoflurane (4% induction, 1.5 – 2% maintenance) and fixed on a stereotaxic frame (Model 942, David Kopf Instruments). After exposing the skull, bregma and lambda were aligned on the same horizontal plane. Small burr holes were then made for viral injections and optic fibers, and additional holes were drilled for anchoring screws. Cre-dependent virus (0.5 µl per side) was injected unilaterally or bilaterally into the PBN (5.0 mm posterior, 1.5 mm lateral, and 3.5 mm ventral to bregma) at a rate of 0.25 µl /min. Microdrives or optic fibers (200 µm diameter, 0.22 numerical aperture) were implanted 0.3 mm dorsal to the virus injection sites and secured with dental cement. Meloxicam (1.5 mg/kg) was administered subcutaneously to alleviate pain and reduce inflammation. Mice were allowed to recover for 2 to 3 weeks before the start of behavioral experiments.

### 30s stimulation of CGRP stimulation

Two weeks after surgery, the optic fibers implanted in the mice were connected to optic cables, and the animals were placed in open arena (30 X 22 X 22 cm). After 2 min of exploration, CGRP neurons were either stimulated or inhibited 4 times at 60-s intervals. For activation, 40 Hz of blue light (473 nm; LaserGlow) was delivered for 30 s. For inactivation, continuous red light (640 nm; LaserGlow) was delivered for 3 s followed by a 1-s ramp down, repeated in cycles until a total duration of 30 s. The light output from the bilateral branching cable was set to 9 ± 0.5 mW. The animals’ behavior was recorded using a camera mounted on the ceiling of the chamber. Freezing and movement velocities were analyzed using video-tracking software (ANY-maze, Stoelting Co.).

### Fear conditioning with chasing robot

Fear conditioning experiment was conducted using a box-shaped robot (15 × 26 × 35 cm) with four high-traction wheels that moved quickly inside a white acrylic track (18 cm width) of a donut-shaped maze (60 cm outer diameter). The speed of the robot was controlled by a Bluetooth-based microcontroller with a custom-written program in Arduino, and the CS (4 kHz; 80 dB; 10 s) was generated by speakers mounted in the front and back of the robot. Habituation and fear responses to the CS were tested before and after conditioning in a rectangular box (30 × 27 × 20 cm). The maze and rectangular box were wiped with 70% ethanol between animals.

On the first day of fear conditioning paradigm, habituation to the CS was performed. Mice were introduced to the rectangular box and allowed a 3-min exploratory period, followed by the presentation of 4 CSs at intervals of 60 s. During this phase, the chasing robot was positioned outside the rectangular box to prevent the animals from seeing it. On day 2, optic fiber cables were attached to the head of each mouse, which were then placed in the donut-shaped maze. After 3 min, mice received 10 associations of a 10-s CS, each co-terminating with 3 s of chasing (speed of 70, 80, or 90 cm/s) with 60-s interval. The robot chased animals at high speeds and posed a physical threat by colliding and pushing them. During robot chasing, CGRP neurons were either activated or inhibited. To optogenetically stimulate, 30 Hz of blue light was delivered for 3 s with the robot. For inhibition, continuous red light was delivered for 3 s followed by a 1-s ramp down with the robot. On day 3, fear response to the CS was measured. Mice were placed in the same rectangular box used on the first day, and after 3 min, the CS was presented alone 4 times at 60-s intervals. During conditioning procedures, the animals’ behavior was recorded using a camera mounted on the ceiling of the chamber. Freezing and movement velocities were analyzed using ANY-maze. Freezing behavior was automatically detected when movement was absent for at least 0.8 s, and a rapid fleeing movement was scored when velocity exceeded 8 cm/s, with a minimum inter-peak interval of 0.6 s. The number of times the mice bumped into the robot was manually scored by an experimenter who was blinded to the group assignments of the animals. We divided each group into 2-3 animal batches and replicated the experiments to confirm the consistency of the results across these batches.

### Fear conditioning with electrical footshock

A standard fear conditioning paradigm with electric footshock was conducted in four identical chambers (21.6 × 17.8 × 12.7 cm; Med Associates) placed inside sound-attenuating boxes. Each chamber was equipped with two speakers on one wall with 24 shock grids on the floor wired to a scrambled shock generator. Tone habituation and retention test of fear memory was tested in a different context where white plastic panels (20 × 16 × 12 cm) were inserted inside the chamber covering the walls and grids. The chamber and inserts were cleaned with 70% ethanol between animals.

After a week of resting period from fear conditioning paradigm with the robot, animals underwent conventional fear conditioning paradigm. On day 1, mice were habituated to a different CS (12 kHz; 80 dB; 10s). The white plastic panels were inserted inside the chamber, and the animals were allowed to freely explore the context for 3 min. The CS was then presented 4 times with an ITI of 60 s. On the next day, after 3 min of free exploration, the animals received 10 CS– US trials, each co-terminating with a 1-s footshock (0.3 mA) with a 60-s ITI. CGRP neurons were either activated or inhibited during footshock delivery. For activation, 30 Hz of blue light was delivered during footshock presentation. For inhibition, continuous red light was delivered for 1 s followed by a 1-s ramp down during footshock presentation. To add a context-specific odor, a petri dish filled with a 1% acetic acid solution was placed under the grid floor. On day 3, fear memory in response to the CS was tested. As on the first day, animals were placed in the chamber with the white plastic panels, and the CS was presented 4 times at 60-s intervals. Animal behavior was recorded during the experiments by a camera installed on the ceiling, and freezing and fleeing responses were analyzed afterward.

### Single-unit recording

A custom-made microdrive containing four tetrodes (20 µm diameter tungsten wire; California Fine Wire) glued to one optic fiber (200-µm core diameter, 0.22 numerical aperture) was used. Tetrode tips were cut to protrude beyond the optic fiber by 400 – 500 µm and were gold-plated to reach impedances of 200-500 kΩ, tested at 1 kHz. After the recovery period from surgery, individual mice were placed in a holding cage, and single-unit activity was monitored using a Cheetah data acquisition system (Digital Lynx SX, Neuralynx). Neural signals were filtered between 0.6 and 6 kHz, digitized at 32 kHz, and amplified 1000 – 8000 times. To identify ChR2-expressing units in the PBN, 10 blue light pulses (473 nm; 5-ms width; 4 – 10 mW/mm^2^ intensity; Laswerglow technologies) were delivered at 30 Hz via the optic fiber. If no light-responsive units were detected, the tetrodes were lowered by 40 – 80 µm increments, up to 160 µm per day. Once light-responsive units were found, behavioral recording sessions began.

During daily recording sessions, spontaneous spikes from PBN neurons were recorded in the home cage for 10 min. Neuronal firing rates were further recorded during fear conditioning sessions over 3 consecutive days. On day 1, mice were first habituated to a tone (10 kHz, 80 dB, 10s duration) as the conditioned stimulus (CS) for 10 times. On day 2, mice underwent 10 exposures to the CS, each co-terminated with an unconditioned aversive stimulus (US) consisting of 3 s of chasing by a robot, with an average inter-trial interval (ITI) of 100 s. On day 3, mice were tested for fear retention with 10 presentations of the CS alone. At the end of each recording session, 10 trains of 10 light pulses (total 100 presentations; 30-s intervals) were delivered to identify ChR2-expressing CGRP neurons in the PBN. The tetrodes were kept in the same location to compare neuronal responses to the CS across three days of conditioning. However, neurons recorded across three days were considered independent units rathern than the same units.

After completing the fear conditioning sessions, neuronal firing rates were recorded in reponse to three different aversive stimuli: pinprick, tail pinch, and robot chasing. Pinprick and tail pinch were given as described previously (Pyeon et al., 2024). For the pinprick, mice were placed in a white cylindrical Plexiglass container (14 cm in diameter, 20 cm in height) with a plastic grid floor, and hind paws were pinpricked with a 26 G syringe needle (approximately 0.5 s duration). For the tail pinch, mice were placed in a rectangular Plexiglass cage (27 × 18 × 8 cm), and the tail was pinched using forceps (1 s duration). For the robot chasing, mice were chased by the same robot used in the conditioning sessions but without the predictive CS. After the daily recording session, all tetrodes were lowered by 40-80 µm to find different light-responsive neurons and the mouse was returned to its home cage.

Neuronal spikes were isolated based on various waveform characteristics using Offline Sorter (Plexon). Stably firing units throughout the behavioral recording session were further analyzed using MATLAB software (MathWorks). To classify CGRP neurons, peri-event time histograms (PETHs; 11.11-ms bins) were constructed around the light presentations. Spike probability and latency were calculated for individual units in response to a total of 100 light pulses, and a cluster analysis was conducted on all units. The cluster with the highest spike probability (> 0.8) and the shortest latency (< 5.5 ms) was identified as CGRP neurons. These neurons also showed higher correlations between spontaneous and light-evoked waveforms, compared with optically insensitive PBN neurons. To further examine responses of CGRP neurons to aversive stmuli, PETHs (50 s-ms bins) were genereated around the time of these aversive stimuli. Firing rates in PETHs were converted to z-scores relative to baseline firing rates observed during 3-s period before each stimulus. Average neuronal responses to aversive stimuli were measured during 0.65-s window from stimulus onset.

### Histology

After completion of all behavioral experiments, mice were anesthetized and transcardially perfused with phosphate-buffered saline (PBS) followed by 4% paraformaldehyde (PFA). Dissected brains were post-fixed overnight in 4% PFA, then cryoprotected in 30% sucrose in PBS at 4°C for 72 h. Brains were frozen and sectioned into 30-µm coronal slices on a cryostat (CM1860, Leica Biosystems). Sections were mounted on microscopic slides and cover-slipped with DAPI Fluoromount-G (Southern Biotech). Using a fluorescence microscope (EVOS M5000, Thermo Fisher Scientific), images were taken to examine recording sites, fiber placements, and fluorescent expression levels.

## Statistical analysis

Statistical analyses were performed using a statistic software package (SPSS version 27.0, IBM SPSS, Armonk, NY). Statistical tests for electrophysiological and behavioral results were assessed with mixed-design ANOVA that contained within-subjects variables (e.g., trials) and between-subjects variables (e.g., group) as well as one-way ANOVA across groups. Once significant interactions were observed, Bonferroni corrections were used for post hoc pairwise comparisons. Two-tailed P values < 0.05 were considered significant. All data were expressed as mean ± SEM.

## Data availability

All data generated or analyzed during this study are included in the manuscript and supporting files.

## Author contribution

G.H.P. and Y.S.J. designed the study. G.H.P., H.C., B.M.C., performed research. G.H.P., B.M.C., and Y.S.J. analyzed the data, and G.H.P., J.-S.C. and Y.S.J. wrote the manuscript.

## Acknowledgements

This work was supported by the National Research Foundation of Korea grant funded by the Korean government (2022M3E5E8017804 to Y.S.J.).

---

**Figure 1-source data 1.** Behavior data for Figure 1.

**Figure 1-source data 2.** Electrophysiology data for Figure 1.

**Figure 2-source data 1.** CGRP stimulation induces an immediate and robust freezing response.

**Figure 3-source data 1.** CGRP stimulation during chasing threat prolongs fleeing responses in aversive memory recall. Behavior data for Figure 3.

**Figure 4-source data 1.** CGRP Stimulation during electric footshock strengthens freezing responses in aversive memory recall.

**Figure 5-source data 1.** Increased robot speed enhances fleeing responses during recall of an aversive memory.

**Figure 5-source data 2.** CGRP neurons are necessary for inducing active defensive behavior under high-speed conditions.

**Figure 1−figure supplement 1.**
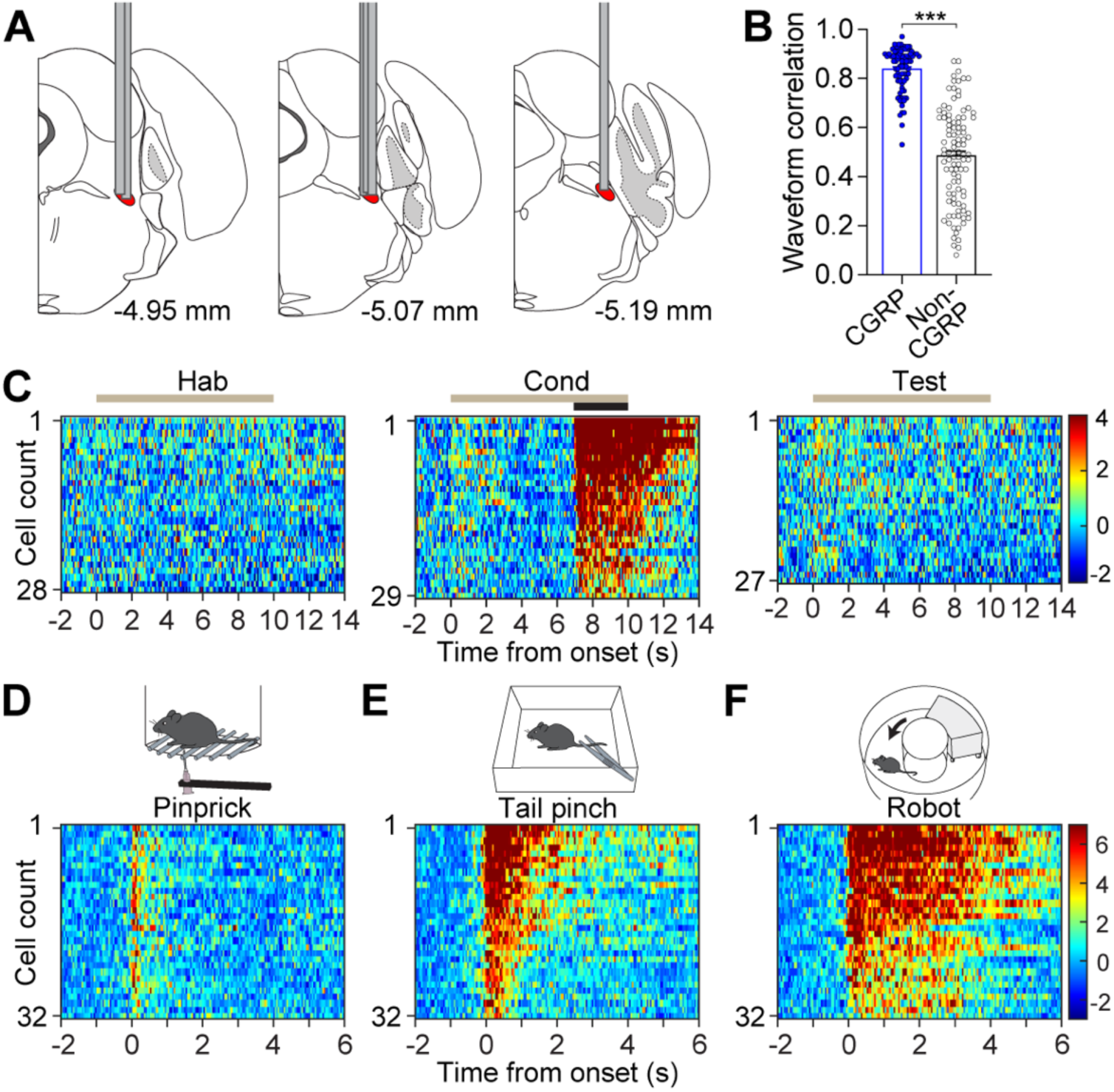
**(A)** Reconstruction of optrode implantation sites along the anterior-posterior extent of the PBN. Each bar indicates an optrode track included for data analysis in the experiment shown in Figure 1 **(B)** Correlations between spontaneous and light-evoked waveforms. CGRP neurons showed higher correlations between spontaneous and light-evoked waveforms compared to non-CGRP neurons (t(181) = 15.16, *p* < 0.001). **(C)** Color-coded, normalized firing rates of CGRP neurons during fear conditioning using the robot. **(D-F)** Color-coded, normalized firing rates of CGRP neurons during (D) pinprick, (E) tail pinch, and (F) chasing. \*\*\**p* < 0.001.

**Figure 3−figure supplement 1.**
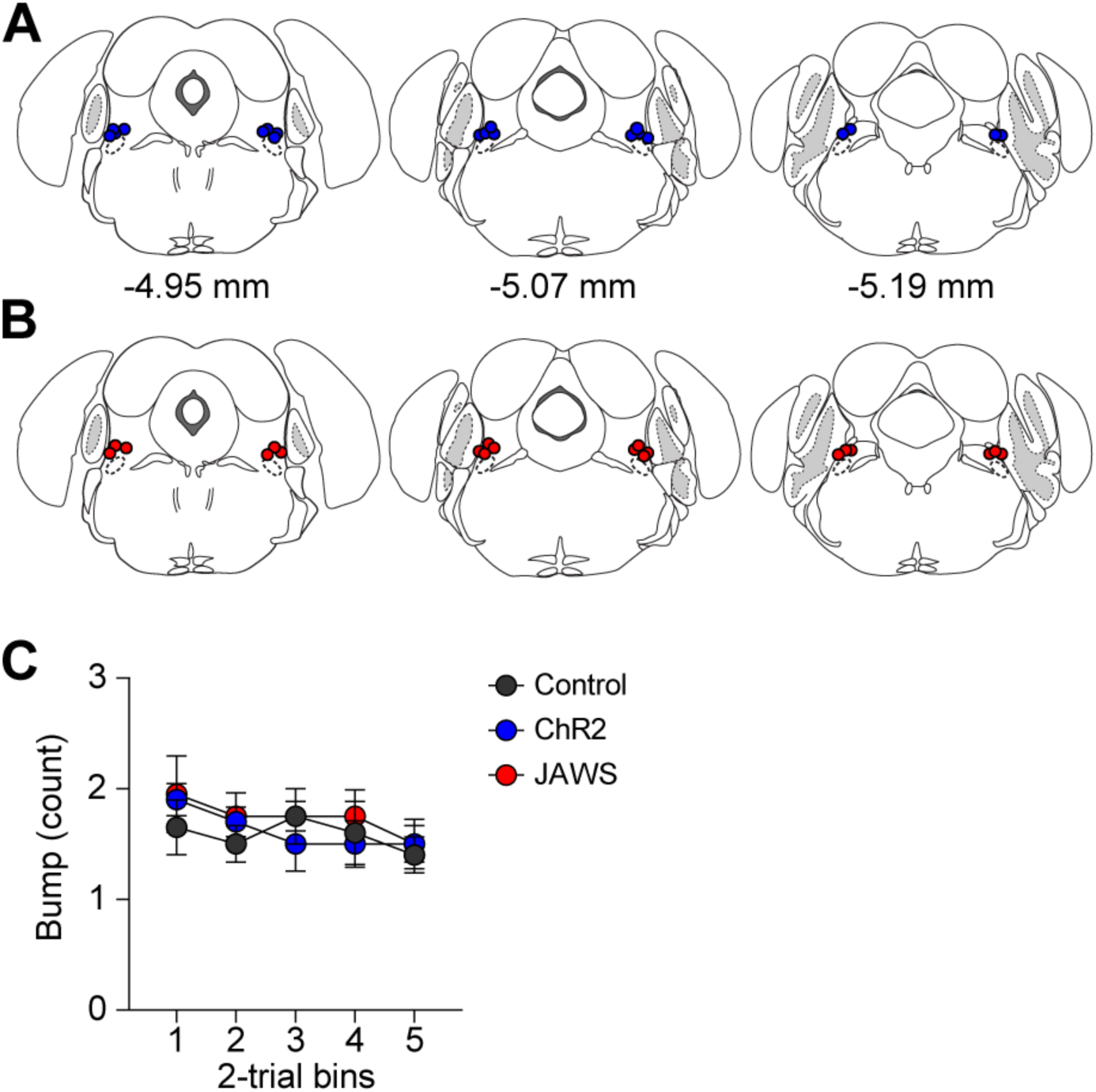
**(A-B)** Anatomical locations of optic fiber tips above the PBN of (A) ChR2 and (B) Jaws groups. **(C)** Number of times bumping with the robot. No group differences were observed throughout the trials across the three groups (a mixed-design ANOVA, F(2, 27) = 0.28, *p* = 0.76).

**Figure 5−figure supplement 1.**
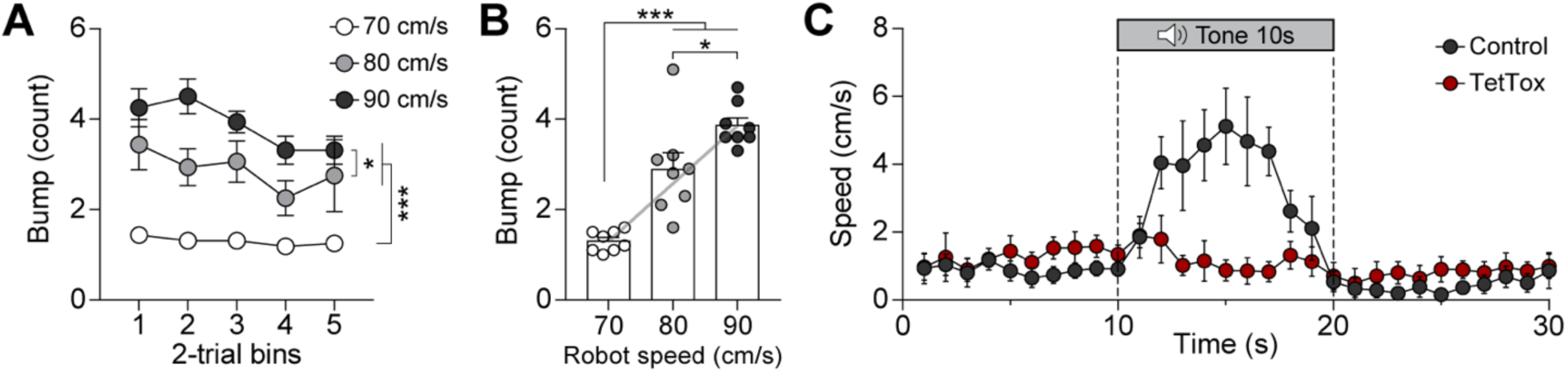
**(A)** Number of times bumping with the robot at different speeds. Bumping occurrence increased with robot speeds (significant group effect in a mixed-design ANOVA, F(2, 21) = 29.44, *p* < 0.001): 80 cm/s resulted in more bumps than 70 cm/s (*p* < 0.001), 90 cm/s resulted in more bumps than 80 cm/s (*p* < 0.05), and 90 cm/s resulted in more bumps than 70 cm/s (*p* < 0.001). **(B)** Average number of times bumping during conditioning. There was a positive correlation between number of bumping and robot speed (gray line; *r* = 0.85, *p* < 0.001). **(C)** Average velocities in response to the CS during the retention test. \**p* < 0.05, \*\*\**p* < 0.001.

**Figure 1−figure supplement 1−source data 1.** Correlations between spontaneous and light-evoked waveforms.

**Figure 3−figure supplement 1−source data 1.** Number of bumping incidents across different groups.

**Figure 5−figure supplement 1−source data 1.** Number of bumping incidents across groups with different robot speeds.

**Figure 5−figure supplement 1−source data 2.** Velocity data for ***Figure5−figure supplement 1E***.

## References

1. Bernard, J., & Besson, J. (1988). Convergence of nociceptive information on the parabrachio-amygdala neurons in the rat. Comptes Rendus de L’academie des sciences. Serie III, Sciences de la vie, 307(19), 841–847.

2. Blanchard, R. J., & Blanchard, D. C. (1969a). Crouching as an index of fear. Journal of comparative and physiological psychology, 67(3), 370.

3. Blanchard, R. J., & Blanchard, D. C. (1969b). Passive and active reactions to fear-eliciting stimuli. Journal of comparative and physiological psychology, 68(1p1), 129.

4. Bolles, R. C. (1970). Species-specific defense reactions and avoidance learning. Psychological review, 77(1), 32.

5. Bolles, R. C., & Collier, A. C. (1976). The effect of predictive cues on freezing in rats. Animal Learning & Behavior, 4(1), 6–8.

6. Borkar, C. D., & Fadok, J. P. (2021). A novel pavlovian fear conditioning paradigm to study freezing and flight behavior. JoVE (Journal of Visualized Experiments)(167), e61536.

7. Bowen, A. J., Chen, J. Y., Huang, Y. W., Baertsch, N. A., Park, S., & Palmiter, R. D. (2020). Dissociable control of unconditioned responses and associative fear learning by parabrachial CGRP neurons. Elife, 9, e59799.

8. Campos, C. A., Bowen, A. J., Roman, C. W., & Palmiter, R. D. (2018). Encoding of danger by parabrachial CGRP neurons. Nature, 555(7698), 617–622.

9. Carter, M. E., Soden, M. E., Zweifel, L. S., & Palmiter, R. D. (2013). Genetic identification of a neural circuit that suppresses appetite. Nature, 503(7474), 111–114.

10. Chen, J. Y., Campos, C. A., Jarvie, B. C., & Palmiter, R. D. (2018). Parabrachial CGRP neurons establish and sustain aversive taste memories. Neuron, 100(4), 891–899. e895.

11. Chiang, M. C., Bowen, A., Schier, L. A., Tupone, D., Uddin, O., & Heinricher, M. M. (2019). Parabrachial complex: a hub for pain and aversion. Journal of Neuroscience, 39(42), 8225–8230.

12. Choi, J.-S., & Kim, J. J. (2010). Amygdala regulates risk of predation in rats foraging in a dynamic fear environment. Proceedings of the National Academy of Sciences, 107(50), 21773–21777.

13. Fadok, J. P., Krabbe, S., Markovic, M., Courtin, J., Xu, C., Massi, L., Botta, P., Bylund, K., Müller, C., & Kovacevic, A. (2017). A competitive inhibitory circuit for selection of active and passive fear responses. Nature, 542(7639), 96–100.

14. Fanselow, M. S. (1980). Conditional and unconditional components of post-shock freezing. The Pavlovian journal of biological science: Official Journal of the Pavlovian, 15(4), 177–182.

15. Fanselow, M. S. (1982). The postshock activity burst. Animal Learning & Behavior, 10(4), 448–454.

16. Fanselow, M. S., & Lester, L. S. (2013). A functional behavioristic approach to aversively motivated behavior:: Predatory imminence as a determinant of the topography of defensive behavior. In Evolution and learning (pp. 185-212). Psychology Press.

17. Fanselow, M. S., & Poulos, A. M. (2005). The neuroscience of mammalian associative learning. Annu. Rev. Psychol., 56, 207–234.

18. Han, S., Soleiman, M. T., Soden, M. E., Zweifel, L. S., & Palmiter, R. D. (2015). Elucidating an affective pain circuit that creates a threat memory. Cell, 162(2), 363–374.

19. Jo, Y. S., Heymann, G., & Zweifel, L. S. (2018). Dopamine neurons reflect the uncertainty in fear generalization. Neuron, 100(4), 916–925. e913.

20. Johansen, J. P., Tarpley, J. W., LeDoux, J. E., & Blair, H. T. (2010). Neural substrates for expectation-modulated fear learning in the amygdala and periaqueductal gray. Nature neuroscience, 13(8), 979–986.

21. Johansen, J. P., Wolff, S. B., Lüthi, A., & LeDoux, J. E. (2012). Controlling the elements: an optogenetic approach to understanding the neural circuits of fear. Biological psychiatry, 71(12), 1053–1060.

22. Juarez, B., Kong, M.-S., Jo, Y. S., Elum, J. E., Yee, J. X., Ng-Evans, S., Cline, M., Hunker, A. C., Quinlan, M. A., & Baird, M. A. (2023). Temporal scaling of dopamine neuron firing and dopamine release by distinct ion channels shape behavior. Science Advances, 9(32), eadg8869.

23. Kang, S. J., Liu, S., Ye, M., Kim, D.-I., Pao, G. M., Copits, B. A., Roberts, B. Z., Lee, K.-F., Bruchas, M. R., & Han, S. (2022). A central alarm system that gates multi-sensory innate threat cues to the amygdala. Cell reports, 40(7).

24. LeDoux, J. E. (2000). Emotion circuits in the brain. Annual review of neuroscience, 23(1), 155–184.

25. Lee, J.-H., Kimm, S., Han, J.-S., & Choi, J.-S. (2018). Chasing as a model of psychogenic stress: characterization of physiological and behavioral responses. Stress, 21(4), 323–332.

26. Lefler, Y., Campagner, D., & Branco, T. (2020). The role of the periaqueductal gray in escape behavior. Current Opinion in Neurobiology, 60, 115–121.

27. Maren, S. (2001). Neurobiology of Pavlovian fear conditioning. Annual review of neuroscience, 24(1), 897–931.

28. Masferrer, M. E., Silva, B. A., Nomoto, K., Lima, S. Q., & Gross, C. T. (2020). Differential encoding of predator fear in the ventromedial hypothalamus and periaqueductal grey. Journal of Neuroscience, 40(48), 9283–9292.

29. Ozawa, T., Ycu, E. A., Kumar, A., Yeh, L.-F., Ahmed, T., Koivumaa, J., & Johansen, J. P. (2017). A feedback neural circuit for calibrating aversive memory strength. Nature neuroscience, 20(1), 90–97.

30. Palmiter, R. D. (2018). The parabrachial nucleus: CGRP neurons function as a general alarm. Trends in neurosciences, 41(5), 280–293.

31. Pyeon, G. H., Kim, J.-H., Choi, J.-S., & Jo, Y. S. (2024). Activation of CGRP neurons in the parabrachial nucleus suppresses addictive behavior. Proceedings of the National Academy of Sciences, 121(24), e2401929121.

32. Pyeon, G. H., Lee, J., Jo, Y. S., & Choi, J.-S. (2023). Conditioned flight response in female rats to naturalistic threat is estrous-cycle dependent. Scientific Reports, 13(1), 20988.

33. Telensky, P., Svoboda, J., Blahna, K., Bureš, J., Kubik, S., & Stuchlik, A. (2011). Functional inactivation of the rat hippocampus disrupts avoidance of a moving object. Proceedings of the National Academy of Sciences, 108(13), 5414–5418.

